# Synergistic effects of multiple pesticides on the flight ability of honey bees (*Apis cerana* F.)

**DOI:** 10.1101/2023.05.22.541595

**Authors:** Changsheng Ma, Xiaoyu Shi, Sihao Chen, Jincai Han, Haodong Bai, Haitao Yu, Zuren Li, Hongmei Li-Byarlay, Lianyang Bai

## Abstract

Pesticides cause risks in the environment for Asian honey bees (*Apis cerana*). Imidacloprid (I), chlorpyrifos (C) and glyphosate (G) are among the most-widely used pesticides in the world. It is not clear on how these pesticides and combination of them affect the flight ability of *A. cerana*. Here we utilized flight mills to show new results that acute treatment of individual pesticides, such as 0.2 ng/bees imidacloprid (20 μL, 10 ng/g), 0.6 ng/bees chlorpyrifos (20 μL, 30 ng/g) and 1.2 ng/bees glyphosate (20 μL, 60 ng/g), had no effect on the flight ability of bees. However, forager bees showed a significantly decrease in the flying duration and flying distance when oral exposed to two or three these pesticides. This evidence indicated that two or three pesticides can produce synergistic changes in the flight ability and behavior of honeybees. Results showed a light on new understandings of complex effects and potential risks of these three pesticide on bee behavior including homing ability and food-collecting ability. Our results are key information to understand new synergistic potential among pesticide formulations and how they impair bee behavior.

**Table of Contents Graphic:** 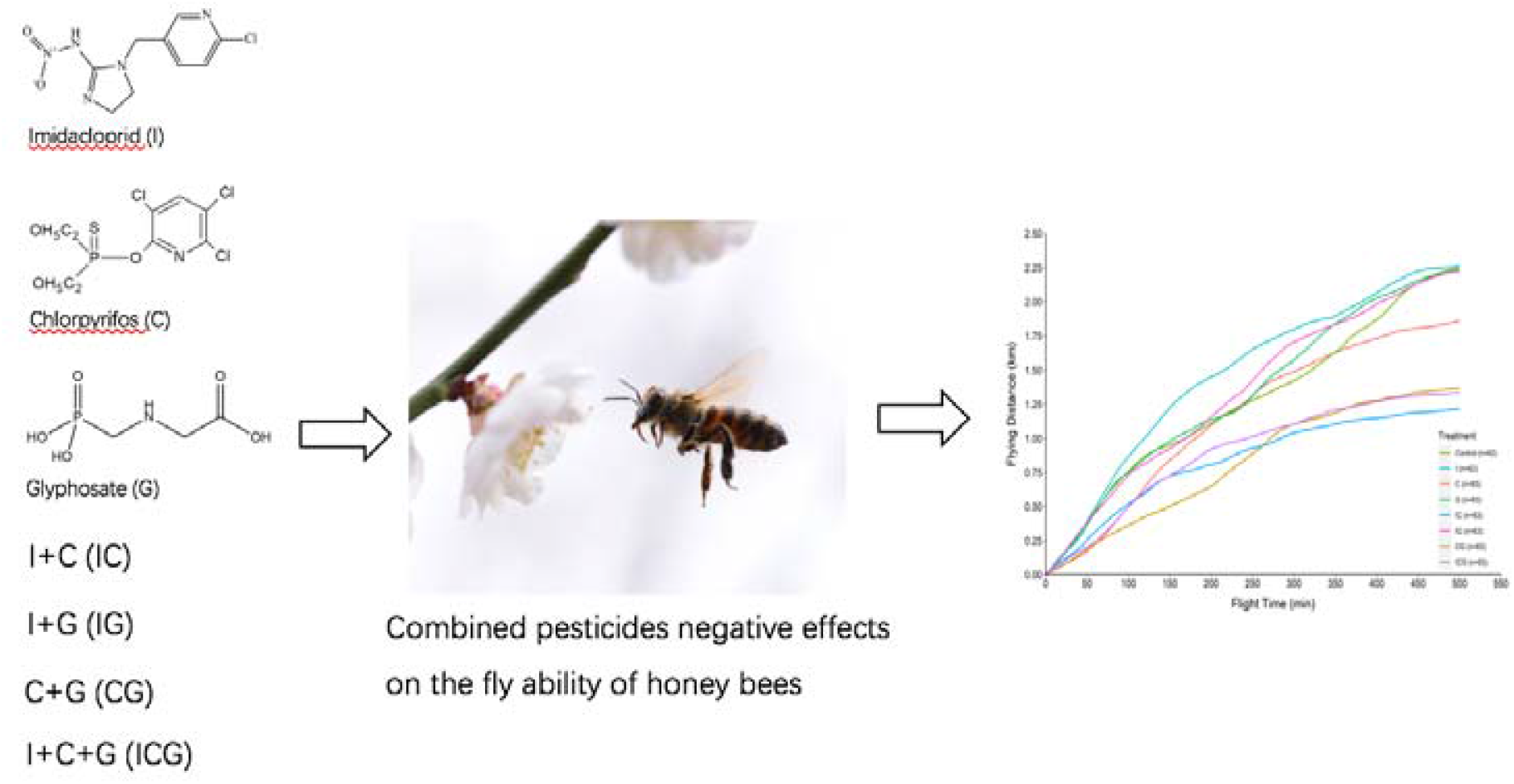

## 1. Introduction

Bees are one of the most important pollinators, providing important pollination services for many crops [1, 2]. Among 20,000 bee species around the world (https://www.discoverlife.org/mp/20q?guide=Apoidea_species), the honey bee as a managed bee, is the most commonly pollinators [3]. However, many factors are threatening the health of managed bees [4, 5]. Widely used pesticides is one of these factors reported by scientific evidence [6-9]. In China, there are mainly two managed honey bee species, *Apis mellifera* and *Apis cerana* [10]. Compared to *A. mellifera*, little research studied the pesticide effects on the health of *A. creana*.

Pesticides are primarily used to protect the value of crops [11]. When the honey bees forage in the field, they are usually exposed to certain levels of pesticides [12, 13]. Residuals of these pesticides are therefore frequently detected in the bees [14, 15], nectar [16, 17] and pollen [18, 19]. Although field residuals of these pesticides did not directly cause the death of honey bees, some evidence showed that these residuals may have a significant effect on the survival of honey bees [20, 21].

Flight ability is essential for the survival of the honey bee colonies because they rely on flying to forage or collect food from many flowers. Impaired flight ability led to less nectar and pollen [20, 22], then reduced the reared number of larva [23], and ultimately caused precocious foraging [24], and increased mortality in bees. Consequently, more younger honey bees went to forage food, which in turn dramatically weakened the colony and accelerated the collapse of the colony [25].

Previous studies have investigated effects of individual neonicotinoid pesticide, and the interactive effect between parasitic mite (*Varroa destructor*) and neonicotinoid pesticide on the flight capability of bees [26-30]. However, to the best of our knowledge, very few study showed the effects of herbicides, organophosphorus insecticides, or their synergistic effects on the flight ability of honey bees. In fact, this is critical to understand the nature of how bees exposed to numerous pesticides when collecting food on different plants in the field [19]. Synergistic effects usually exacerbate individual effect, and are common in the fields compared to laboratory conditions [31]. Other studies showed the combinatorial effects of pesticide exposure on bees but flying behavior was not investigated previously [32].

The aim of this study is to investigate the influences of acute oral exposure of three widely used pesticides, an insecticide called imidacloprid, an organic phosphate chemical called chlorpyrifos, and a widely used herbicide called glyphosate on the flight ability of honey bees. Our research questions include what the impact of single imidacloprid, chlorpyrifos and glyphosate independently or combined effects on the flight ability of *A. cerana* is.

## 2. Material and Methods

### (a) Pesticide preparation

All pesticides were obtained from Aladdin, Inc. The names, CAS numbers, and purity information of each pesticide were as follows: (i) imidacloprid: CAS No. 138261-41-3, purity of 98.5%; (ii) chlorpyrifos: CAS No. 2921-88-2, purity of 99%; and (iii) glyphosate: CAS No. 1071-83-6, purity of 99.5%. Imidacloprid and chlorpyrifos were dissolved in acetone to make a stock solution, while glyphosate was dissolved in water to make a stock solution. These stock solutions were then diluted with 50% (w/w) sucrose solution to make a concentration of 10 ng/g imidacloprid (I), 30 ng/g chlorpyrifos (C), 60 ng/g glyphosate (G), I + C (IC), I + G (IG), C + G (CG), I + C + G (ICG) contaminated sucrose solution. A control sucrose solution without pesticides was also prepared. Previous studies suggested that these concentration was widely found in nectar, pollen and beebread collected from the fields [17, 18, 33, 34].

### (b) Honey bee preparation

Three healthy beehive colonies were reared at Hunan Academy of Agriculture Sciences (HAAS, 113.093483 E, 28.206193 N, Changsha, China), where the study was conducted from May 2020 to July 2020. First, we closed the entrance of the beehive and then collected bees who carrying pollen to ensure that the collected bees share a similar age and flight ability. Subsequently, bees were minimally chilled by ice, and then connected with a 1-cm-long hollow Teflon tube glued to the top of the thorax (see Ma *et al*. [27]), and ultimately fed 20 μL of either the 50% (w/w) sucrose solution containing I, C, G, IC, IG, CG, ICG, and the control solution.

### (c) Flight ability assessment

After feeding treatments, bees were put on 24 flight mills (Jiaduo Industry & Trade Co., Ltd, Hebei, China, FXM-2) for about six hours until most of bees flying distance no longer increased (Fig. 1). During the flight of the bee, since bees did not fly continuously instead intermittently, so we counted the total number of flights they took off during the whole experiment. Among these number of flights, we computed mean flying distance for each flight, mean flying duration for each flight, maximum flying distance among all flights, max flying duration among all flights, and max flying velocity among all flights, total flying distance, total flying duration, and flying velocity of each bee.

**Figure 1.**
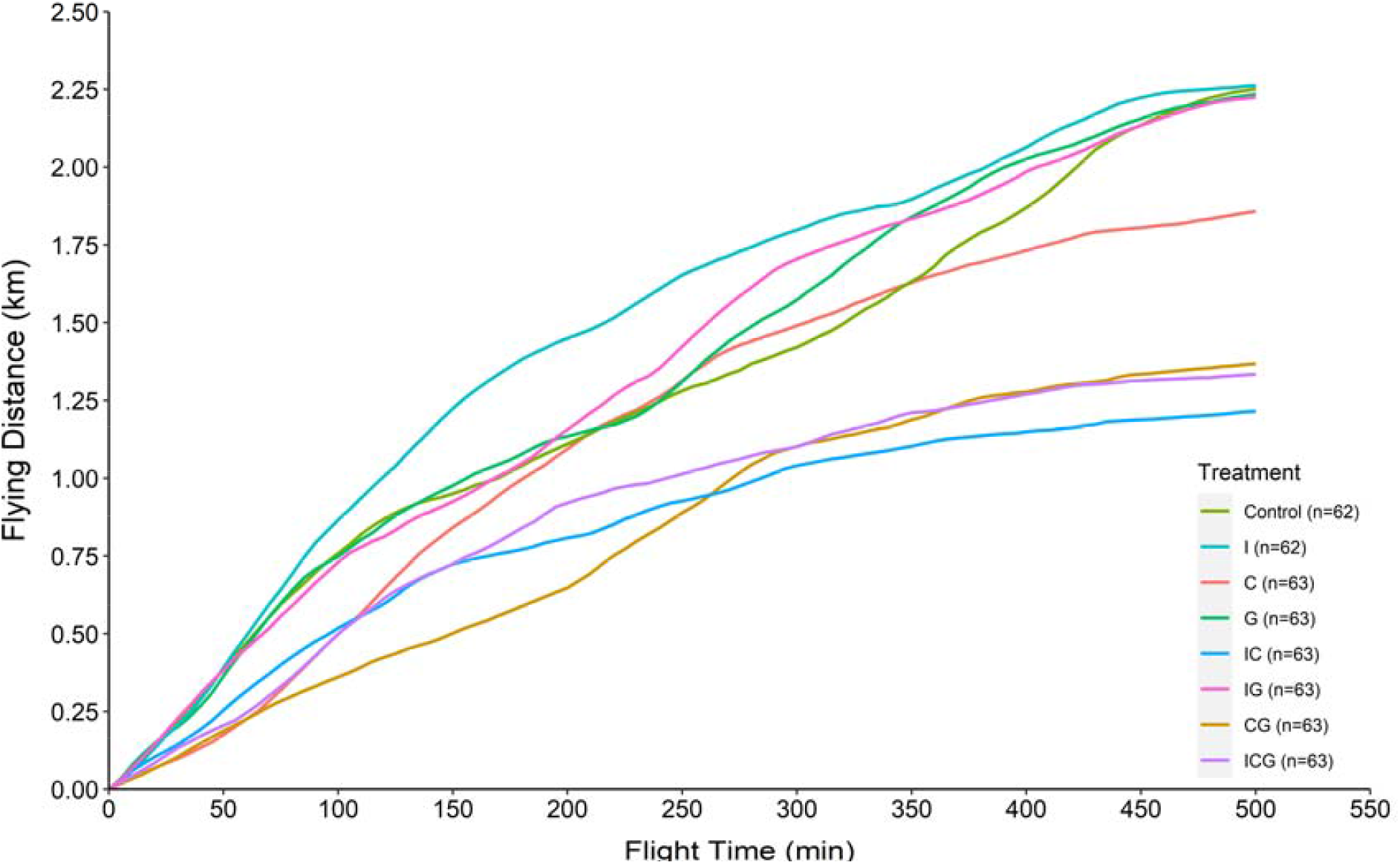
Comparison of the effect of the eight different treatment groups on cumulative flying distance with time to show the effects of exposure to I, C, G or combinations on flight distance of foragers. The figure legend indicated the color and sample size of each group in the graph.

Twenty-four foragers were selected randomly in a colony and then separated into eight different treatment groups (e.g., 7 pesticide treatments and 1 control treatment). This process repeated seven times. Three colonies were used. In total, each treatment consisted of 62 or 63 foragers, and 502 foragers were used in this experiment. All flight mills were located in a climate-regulated room with constant air temperature (25°C), light (5700 lux).

### (d) Statistical analyses

The effects of the eight different treatment groups on the flight ability of bees are tested using One-way ANOVA analyses in GraphPad Prism (GraphPad Prism 8.0, United States). Data were first analyzed using the Shapiro-Wilk normality test to verify the normal distribution assumption, and then using post-hoc Tukey’s test to compare between any two treatments. *P* < 0.05 was considered statistically significant.

## 3. Results

To test the flight ability, we measured the (i) total number of flights, (ii) mean flying distance for each flight, (iii) mean flying duration for each flight, (iv) maximum flying distance among all flights, (v) max flying duration among all flights, (vi) max flying velocity among all flights, (vii) mean flying velocity, (viii) total flying distance, and (ix) total flying duration for each bee under different treatment groups.

Results from the flight mill showed that no significant difference was found between the control and pesticide-treated bees for the total number of flights (*P >* 0.05 in all cases, Fig. 2a), mean flying duration for each flight (*P >* 0.05 in all cases, Fig. 2b), mean flying distance for each flight (*P >* 0.05 in all cases, Fig. 2c), max velocity among all flights (*P >* 0.05 in all cases, Fig. 3a), and mean velocity (*P >* 0.05 in all cases, Fig. 4a). However, bees treated with IC (Df = 494, Tukey HSD Q statistic = 4.30, *P* = 0.05, Fig. 3b), CG (Df = 494, Tukey HSD Q statistic = 4.47, *P* = 0.036, Fig. 3b), and ICG (Df = 494, Tukey HSD Q statistic = 4.39, *P* = 0.042, Fig. 3b) had a significantly lower maximum flying duration among all flights, and maximum flying distance among all flights (Df = 493, Tukey HSD Q statistic = 4.39, *P* = 0.042 for IC; Df = 493, Tukey HSD Q statistic = 4.34, *P* = 0.046 for CG; Df = 493, Tukey HSD Q statistic = 4.35, *P* = 0.046 for ICG, Fig. 3c) than the control group. In addition, our results also showed that there was a significantly higher total flying duration (Df = 494, Tukey HSD Q statistic = 4.44, *P =* 0.038, Fig. 4b) and flying distance (Df = 493, Tukey HSD Q statistic = 4.71, *P =* 0.021, Fig. 4c) for bees treated with IC in comparison with control.

**Figure 2.**
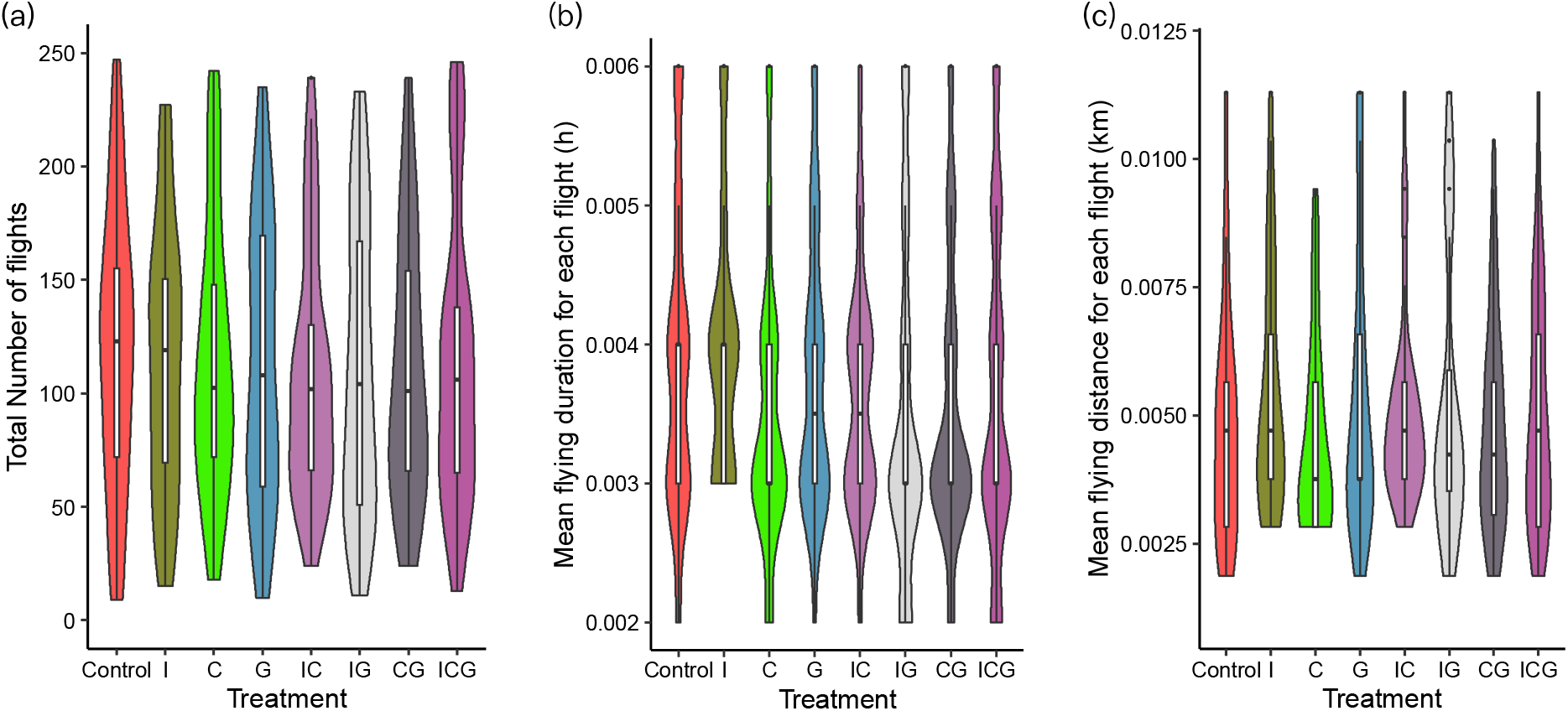
The effects of three pesticides on flights of foragers showing in violin plots and box plots in the middle. We recorded and compared the effect of the individual pesticide and combinations (total of eight treatment groups) on (a) total number of flights, (b) mean flying duration of each flight, and (c) mean flying distance of each flight. The post-hoc Tukey test showed no difference among groups. (N_control_ = 62, N_I_=62, N_C_=63, N_G_ = 63, N_IC_=63, N_IG_ = 63, N_CG_ = 63, N_ICG_ = 63).

**Figure 3.**
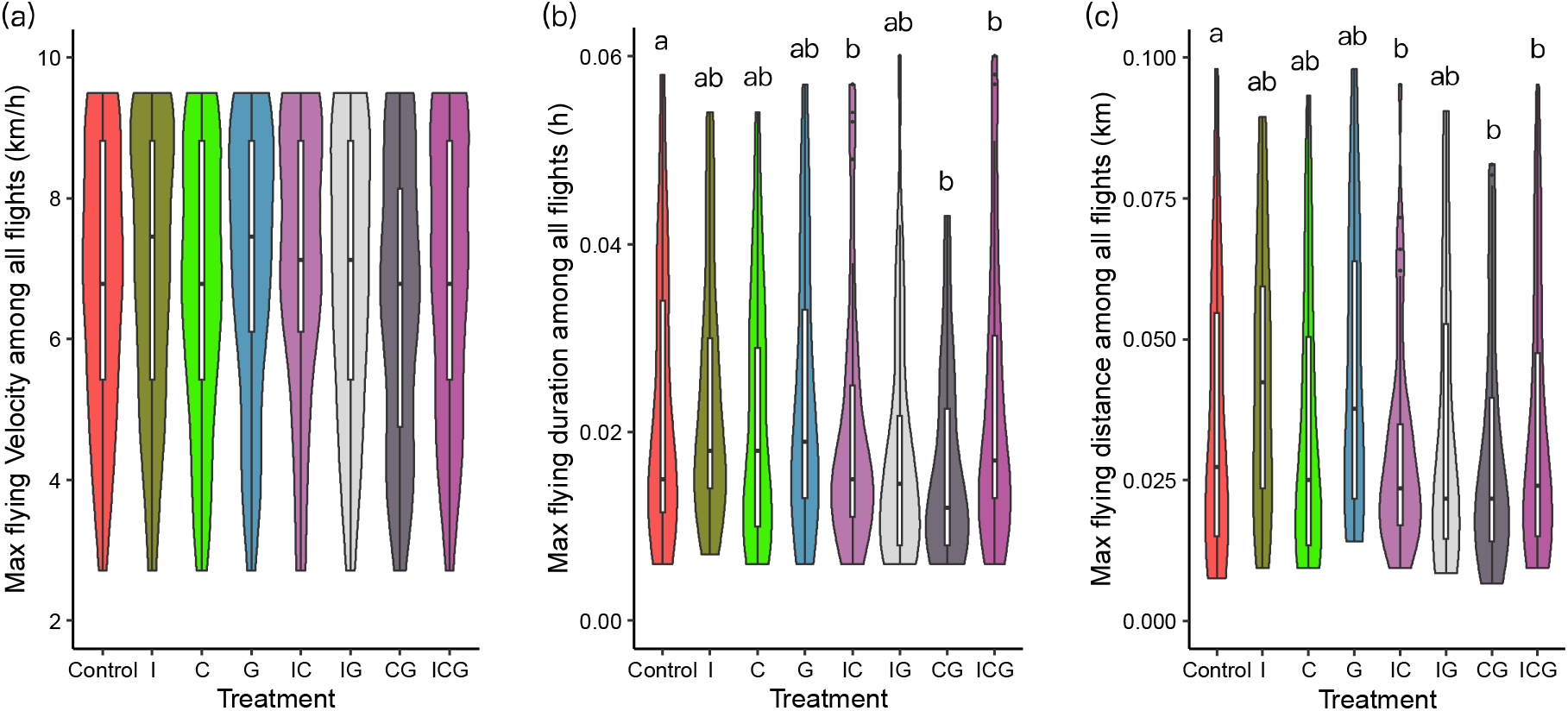
Comparison of the effect of three pesticides on fly velocity, flying duration, and max flying distance in foragers. We recorded and compared the effect of the individual pesticide and combinations (total of eight treatment groups) on (a) max flying velocity, (b) mean flying duration, and (c) max flying distance of each flight. We listed the value of each group in a violin plot with the median in the middle. There was no significant difference in (a) max flying velocity. But there are significant difference among treatments in (b) and (c). Different letters indicate significant differences based on the post-hoc Tukey test. (N_control_ = 62, N_I_ = 62, N_C_ = 63, N_G_ = 63, N_IC_ = 63, N_IG_ = 63, N_CG_ = 63, N_ICG_ = 63).

**Figure 4.**
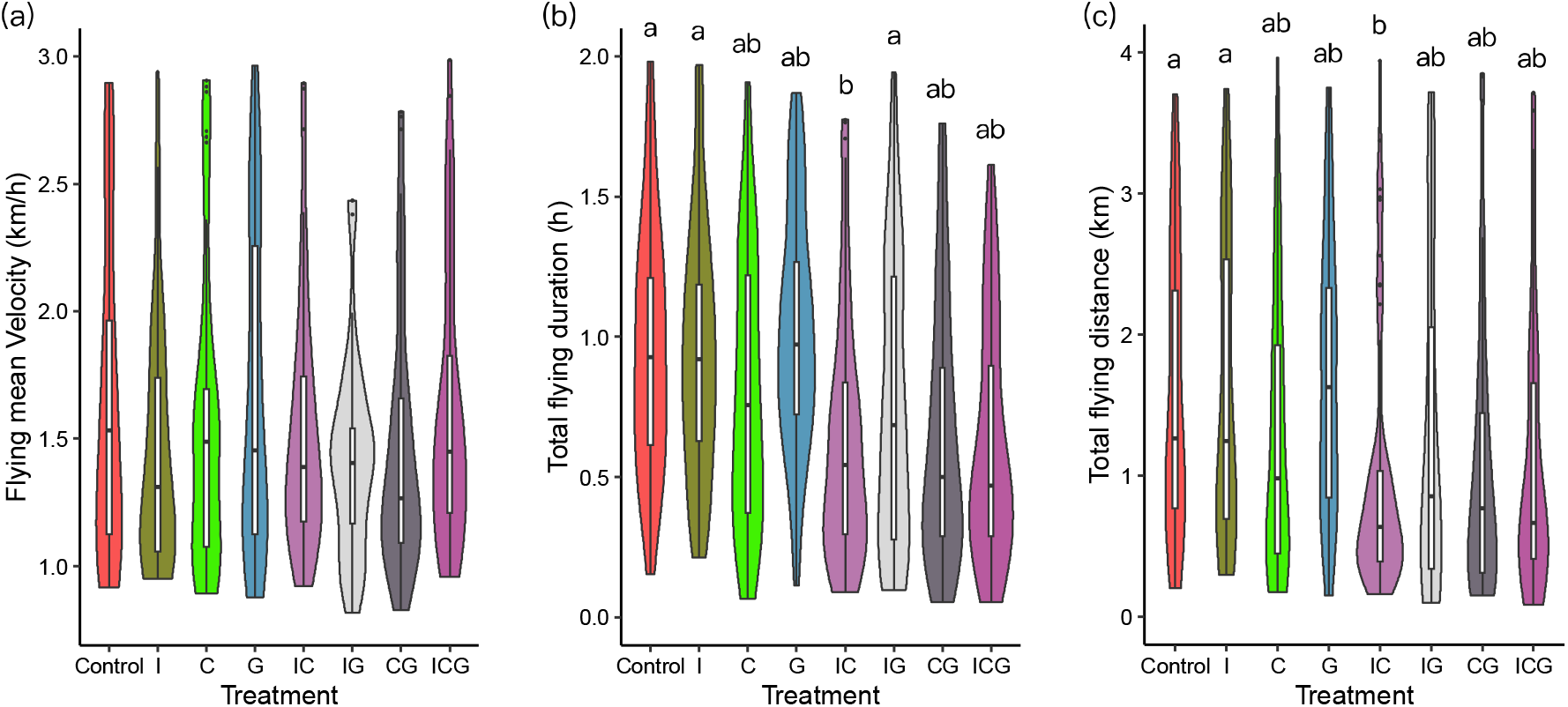
Comparison of the effect of three pesticides on (a) flying mean velocity, (b) flying duration, and (c) flying distance of foragers. We recorded and compared the effect of the individual pesticide and combinations (total of eight treatment groups) on (a) flying mean velocity, (b) flying duration, and (c) flying distance. We listed the value of each group in a violin plot with the median in the middle. There is no significant difference in (a) flying mean velocity, but significant difference among groups in (b) total flying duration and (c) total flying distance. Different letters indicate significant differences based on the post-hoc Tukey test. (N_control_ = 62, N_I_ = 62, N_C_ = 63, N_G_ = 63, N_IC_ = 63, N_IG_ = 63, N_CG_ = 63, N_ICG_= 63.

In comparison, bees treated with I (Df = 494, Tukey HSD Q statistic = 5.05, *P* = 0.0093, Fig. 4b) and IG (Df = 494, Tukey HSD Q statistic = 4.34, *P =* 0.047 Fig. 4b) has a significantly higher total flying duration than bees treated with IC. Moreover, bees treated with I had a significantly higher flying distance then IC treatment (Df = 493, Tukey HSD Q statistic = 4.39, *P* = 0.042, Fig. 4c).

## 4. Discussion

We show in this study for the first time that flying distance and flying duration were all reduced dramatically when bees were exposed to combined pesticides with field-realistic doses. Our results showed that *A. cerana* can fly up to 2.69 km without exposure to pesticides, which is similar to *A. mellifera* that fly up to 1.8 km [35] or 2.1 km [26]. However, the flying distance of *A. cerana* increased about 92% compared to one previous study in *A. cerana* [27]. This may be because the higher sucrose concentrations (e.g., 20 μL 50% w/w sucrose solution vs 20 μL 40% w/w sucrose solution) and higher light intensity (i.e., 5700 lux vs 546 lux) allow bees to have higher flight distance [28].

Our results also suggested that exposure to any of single pesticide treatment, such as I treatment (20 μL of 10 ng/g imidacloprid, 0.02 ng/bee), C treatment (20 μL of 30 ng/g chlorpyrifos, 0.06 ng/bee), and G treatment (20 μL of 60 ng/g glyphosate, 0.12 ng/bee), had no effect on flight ability of bees. Another study [27] showed that a dose of 20 μL of 10 ng/g thiamethoxam (TMX) (0.02 ng/bee) can significantly affected bee flight time, flight speed and flight distance. One explanation is that TMX is more toxic to bees than imidacloprid (LC50 _TMX_ = 8.23 ± 0.56 μg/g vs LC50 _I_ = 24.09 ± 1.19 μg/g) [36, 37].

In addition, the latest study found that chronic exposure to glyphosate could impair the ability of bumblebees to regulate temperature [38]. Negative temperature regulation may result in a decrease in the temperature of skeletal muscles, which in turn impairs the flight ability of bees [39]. However, our results demonstrated that acute exposure to glyphosate did not significantly affect the flight ability of bees, which might be due to the different sensitivity to pesticides between bumblebees and honeybees [40], or to different exposing styles (i.e., chronic exposed vs acute chronic).

These results do not mean that bees are safe and unaffected when exposed to field level imidacloprid and chlorpyrifos. On the contrary, bees treated with the combined 10 ng/g imidacloprid and 30 ng/g chlorpyrifos (IC) solution had a significantly lower flying duration and flying distance than the control group. In particular, bees exposed to IC could only fly 56.14% of the total duration, and 52.04% of total distance of non-exposed bees (e.g., 0.64 h and 1.40 km for bees exposed to IC, vs 1.4 h and 2.69 km for control bees). Furthermore, we found that CG and ICG also reduced the maximum flying duration among all flights and maximum flying distance among all flights. This might be because two or three pesticides can be synergistic, in which the effect of combined stressors is significantly higher than single stressor [41, 42]. Most importantly, bees usually fly a distance of 5.5 km for foraging [43], and therefore are frequently exposed to a number of pesticides [12], such as imidacloprid and chlorpyrifos often appearing simultaneously in the food of bees in field settings [17, 18]. However, most studies only assessed the effects of single pesticide on the health of bees. Our results provided the detrimental synergistic effects from combined pesticides to bee behavior, which highlighted more severe risks than single pesticides. This is true for the flight ability of bees in the real environments.

Many previous research [44, 45] showed that multiple pesticides in the field can cause severe disorders and change in behavior of worker honey bees. Our data is the new additional evidence to show the similar detrimental effects of pesticides in honey bee behavior. Exposure of three pesticides may affect the gene expression of the metabolism pathways for energy as a new research indicated [46]. More studies are needed to elucidate the molecular mechanism of how the synergistic effects of multiple pesticides affect the flight ability, the muscle, and the behavior of foraging in worker bees.

In conclusion, we used flight mills to study the impact of acute oral exposure to I, C, G, IC, IG, CG, ICG on the flight ability of *A. cerana*. We concluded that single pesticide treatment had no effect on the ability of bees. However, when exposed to two or three pesticides, the flying duration and flying distance of bees was significantly reduced. Our results show that exposure to combined pesticides can have synergistic effects in the flight duration and distance of bees. It explains the observed phenomenon of decreased homing ability of bees in the fields, and also colony collapses and reduced pollination services. Our research adds new knowledge when trying to fully understand the costs associated with multiple pesticides in the fields, showing that both the potential synergistic effects and the proper management of pesticides usage need to be considered.

## Authors’ contributions

C.M., Z.L. and L.B. conceived and designed the experiments. C.M., J.H., H.B., performed the experiments. C.M., H.L-B. and X.S. analyzed the data. C.M., X.S., H.L-B., and S.C. wrote the manuscript, Y.H. and other coauthors provided comments and revision.

## Conflicts of interest

The authors declare no competing financial interests.

## Funding

This research was partially funded by the Natural Science Foundation of China (32172433), Natural Science Foundation of Hunan Province (2021JJ20034), the China Agriculture Research System of MOF and MARA (CARS-16-E19), the Hunan Agriculture Research System (2022-31), and Scientific-Innovative of Hunan Agricultural Sciences and Technology (2022CX70, 2021CX42 and 2022CX11).

## Acknowledgments

We would like to thank Xiaoling Wen for her assistant on preparing samples. We also thank Yi Zou, Shudong Luo for their assistance on the resources and contributed to this research.

## Notes

### Competing Interest Statement

The authors have declared no competing interest.

